# Residual Microglia Following Short-term PLX5622 Treatment in 5xFAD Mice Exhibit Diminished NLRP3 Inflammasome and mTOR Signaling, and Enhanced Autophagy

**DOI:** 10.1101/2024.07.11.603157

**Authors:** Maheedhar Kodali, Leelavathi N. Madhu, Yogish Somayaji, Sahithi Attaluri, Charles Huard, Prashanta Kumar Panda, Goutham Shankar, Shama Rao, Bing Shuai, Jenny J. Gonzalez, Chris Oake, Catherine Hering, Roshni Sara Babu, Sanya Kotian, Ashok K. Shetty

## Abstract

Chronic neuroinflammation represents a prominent hallmark of Alzheimer’s disease (AD). While moderately activated microglia are pivotal in clearing amyloid beta (Aβ), hyperactivated microglia perpetuate neuroinflammation. Prior investigations have indicated that the elimination of ∼80% of microglia through a month-long inhibition of the colony-stimulating factor 1 receptor (CSF1R) during the advanced stage of neuroinflammation in 5xFamilial AD (5xFAD) mice mitigates synapse loss and neurodegeneration without impacting Aβ levels. Furthermore, prolonged CSF1R inhibition diminished the development of parenchymal plaques. Nonetheless, the immediate effects of short-term CSF1R inhibition during the early stages of neuroinflammation on residual microglial phenotype or metabolic fitness are unknown. Therefore, we investigated the effects of 10-day CSF1R inhibition in three-month-old female 5xFAD mice, a stage characterized by the onset of neuroinflammation and minimal Aβ plaques. We observed ∼65% microglia depletion in the hippocampus and cerebral cortex. The leftover microglia demonstrated a noninflammatory phenotype, with highly branched and ramified processes and reduced NOD-, LRR-, and pyrin domain-containing protein 3 (NLRP3) inflammasome complexes. Moreover, plaque-associated microglia were reduced in number with diminished Clec7a (dectin-1) expression. Additionally, both microglia and neurons displayed reduced mechanistic target of rapamycin (mTOR) signaling and autophagy. Biochemical assays validated the inhibition of NLRP3 inflammasome activation, decreased mTOR signaling, and enhanced autophagy. However, short-term CSF1R inhibition did not influence Aβ plaques, soluble Aβ-42 levels, or hippocampal neurogenesis. Thus, short-term CSF1R inhibition during the early stages of neuroinflammation in 5xFAD mice promotes the retention of homeostatic microglia with diminished inflammasome activation and mTOR signaling, alongside increased autophagy.

## Introduction

Alzheimer’s disease (AD) pathology is characterized by persistent neuroinflammation, the presence of extracellular amyloid-beta (Aβ) plaques, and intracellular neurofibrillary tangles (Hanslik and Ulland, 2020). Activated microglia and reactive astrocytes are well-recognized contributors to the chronic neuroinflammatory state observed in various neurological and neurodegenerative conditions, including AD (Gomez-Nicola et al., 2015; Simon et al., 2019). Microglial activation, in response to pathogenic invasion or brain injury, is initially beneficial, facilitating the clearance of pathogens or cellular debris through the release of proinflammatory cytokines and engagement in phagocytosis, thereby promoting tissue repair (Olmos-Alonso et al., 2016; Cai et al., 2014). However, sustained microglial activation, beyond the removal of pathogens or cellular debris, instigates chronic neuroinflammatory cascades characterized by excessive production of proinflammatory cytokines and reactive oxygen species, culminating in progressive neurodegeneration (Lyman et al., 2014; Calsolaro et al., 2016). In conditions such as AD, microglial activation occurs in response to the extracellular accumulation of Aβ (Olmos-Alonso et al., 2016). Such disease-associated microglia (DAM) perform phagocytic remodeling of Aβ plaques and inhibit plaque extension via barrier formation around plaques. However, they also contribute to disease progression by releasing multiple detrimental proinflammatory cytokines (Condello et al., 2015; Song et al., 2018). The continuous release of elevated concentrations of proinflammatory cytokines such as interleukin-1 beta (IL-1β) and IL-18 occurs as a result of increased activation of NOD, LRR, and pyrin-domain containing 3 (NLRP3) inflammasomes in activated microglia, which can initiate downstream inflammatory cascades, including the chronic activation of p38/MAPK signaling, resulting in the continuous release of multiple other proinflammatory cytokines (Kheiri et al., 2018) and changes in astrocyte morphology and function and complement activation (Tenner et al., 2020).

Activated microglia also impact the function of neurons and promote synapse loss and neurodegeneration (McFarland et al., 2022). Consequently, numerous studies in AD models have focused on evaluating the implications of microglial elimination at various disease stages. In these investigations, partial or near-total elimination of parenchymal microglia was achieved by inhibiting the colony-stimulating factor 1 receptor (CSF1R). This receptor is prominently expressed in myeloid cells, including microglia, and is pivotal for microglial survival and proliferation (Pixley et al., 2004; Stanley et al., 2014; Waisman et al., 2015). Studies in disease models have revealed that pharmaceutical inhibition of CSF1R induces apoptotic microglial death, resulting in a reduction in the overall microglial population and proinflammatory cytokine levels within a few days (Elmore et al., 2014; Spangenberg et al., 2019; Mancuso et al., 2019). Furthermore, cessation of CSF1R inhibition prompts the proliferation of residual microglia, culminating in the repopulation of microglia to nearly normal levels in the brain within weeks (Elmore et al., 2014; 2015). Hence, selective elimination and restoration of the microglial population offer valuable insights into the role of microglia in the pathogenesis of neurological and neurodegenerative conditions.

A previous study employing a month-long CSF1R inhibition in 10-month-old 5x familial AD (5xFAD) mice (representing an advanced AD stage) reported that ∼80% microglia depletion does not result in reduced Aβ levels or plaques. However, this intervention effectively limited synapse loss and neurodegeneration (Spangenberg et al., 2016). Furthermore, the depletion of ∼95% of microglia either before the onset of neuroinflammation in AD (1.5-month-old 5xFAD mice) or in the late stage of AD (14-month-old 5xFAD mice) did not produce any discernible effect on Aβ plaques (Spangenberg et al., 2016). However, studies employing CSF1R inhibition for extended periods in 5xFAD mice reported diminished parenchymal plaque development (Spangenberg et al., 2019; Son et al., 2020). Moreover, a two-week CSF1R inhibition resulting in approximately 50% depletion of microglia in 22-month-old 3xTg mice and ∼65% depletion in 14-month-old APP/PS1 mice did not lead to a reduction in Aβ-plaques or neurite damage, although the repopulated microglia exhibited trends of reduced activation (Karaahmet et al., 2022). Other studies have suggested that eliminating microglia in AD models enhanced neural circuit connectivity and activity (Liu et al., 2021) and mitigated tau pathology and neuronal atrophy (Lodder et al., 2021).

To date, no studies have examined the immediate effects of short-term CSF1R inhibition in the early stages of AD on residual microglial morphology or their metabolic fitness. It remains unclear whether short-term CSF1R inhibition in the early stages of AD would predominantly eliminate activated microglia with NLRP3 inflammasome complexes or homeostatic microglia. Our investigation focused on the impact of 10-day CSF1R inhibition on residual microglia in three-month-old female 5xFAD mice, a critical stage when neuroinflammation has commenced but Aβ plaques are minimal. We sought to answer the question: Would ten days of CSF1R inhibition in the early stage of amyloidosis significantly deplete activated microglia in 5xFAD mouse brain, resulting in residual microglia exhibiting a noninflammatory phenotype characterized by highly ramified processes, reduced NLRP3 inflammasome complexes and mTOR signaling, and improved metabolic fitness? If so, how would it impact the extent of NLRP3 inflammasome activation, neuronal mTOR signaling and autophagy, soluble Aβ-42, and Aβ plaque levels in the hippocampus and the cerebral cortex? Furthermore, our study included an evaluation of whether the altered microenvironment following the 10-day CSFIR inhibition would influence the status of hippocampal neurogenesis.

## Materials and Methods

### Animals

The study employed three-month-old female 5xFAD mice with five human familial AD mutations driven by the mouse Thy1 promoter and age-matched wild-type mice (B6SJL) (Oakley et al., 2006). The AD mice were bred in-house and maintained in the vivarium of Texas A&M University. The 5xFAD transgenic male mice and B6SJLF1/J female mice employed for breeding were procured from Jackson Laboratories (Cat No: 34840-JAX and 100012-JAX, Bar Harbor, ME, USA). Animals were housed in an environmentally controlled room with a 12:12-h light: dark cycle and were given food and water ad libitum. The Animal Care and Use Committee of Texas A&M University approved all experimental procedures performed in the study.

### PLX5622 Treatment and Study Design

Three-month-old 5xFAD female mice were randomly assigned to 2 groups: one group received AIN-76A standard rodent chow containing PLX5622 (n=12, AD+PLX group), and another group received AIN-76A chow without PLX (n=12, AD group) for ten days. The drug PLX5622 was purchased from Medkoo Biosciences (Morrisville, NC, USA), which was formulated into AIN-76A standard chow at the dose of 1200 ppm by Research Diets Inc. (New Brunswick, NJ, USA). Age-matched wild-type mice that received standard lab chow in parallel served as naïve controls (n=12, Naïve group). At the end of the 10-day diet, as described above, 50% of mice from each group (n=6/group) were deeply anesthetized, perfused with 4% paraformaldehyde, and the brain tissues were postfixed in 4% paraformaldehyde, and processed for cryostat sectioning. Fresh brain tissues were harvested following deep anesthesia and decapitation from the remaining 50% of mice in each group (n=6/group). The tissues were snap-frozen and stored at −80°C until processed for biochemical assays.

### Tissue processing and immunohistochemistry

The brain tissues were processed and sectioned using a cryostat as described elsewhere (Rao et al., 2008; Hattiangady and Shetty 2011; Kodali et al., 2018). Thirty-micrometer thick coronal sections through the entire brain were collected in 24-well plates containing phosphate buffer (PB) and stored in a cryobuffer at −20°C until processed for immunohistochemistry. Serial sections (every 20th through the hippocampus and the cerebral cortex) were processed for immunohistochemical detection of the Ionized calcium-binding adaptor molecule 1 (IBA-1, a marker of microglia), glial fibrillary acidic protein (GFAP, a marker of astrocytes) and Aβ42 (a marker of Aβ plaques). Furthermore, every 15th section through the hippocampus was employed to visualize doublecortin (DCX) positive newly born neurons. The methods employed for immunohistochemistry are described in our previous reports (Kodali et al., 2018; Madhu et al., 2019, 2021). The primary antibodies employed in the study include goat anti-IBA-1 (1:1000, Abcam, Cambridge, MA, USA); rabbit anti-GFAP (1:2000, DAKO, Santa Carla, CA, USA); rabbit anti-Aβ42 (1:500, Invitrogen, Grand Island, NY, USA); and rabbit anti-DCX (1:1000, Synaptic systems, Göttingen, Germany). The secondary antibodies comprised biotinylated anti-rabbit or anti-goat IgG (Vector Labs, Burlingame, CA, USA). The peroxidase reaction was developed using Vector SG (Vector Labs) as the chromogen. After a thorough wash, the sections were mounted on subbed slides, counterstained with nuclear fast red (Vector Labs), coverslipped, and observed under a Nikon E600 microscope.

### Preparation of brain tissue lysates for biochemical assays

Hippocampal and cerebral cortex tissues from naïve control, AD, and AD-PLX groups were micro-dissected. The tissues were lysed through sonication in a tissue extraction reagent (Invitrogen) containing protease-phosphatase inhibitor (1:100 dilution, ThermoFischer Scientific, Waltham, MA) for 15-20 seconds at 4°C. The resulting solution was centrifuged at 4°C for 10 minutes (15000g), and the supernatant was aliquoted and stored at −80° C until further use. The lysates were used to measure the concentration of multiple markers.

### Dual or triple immunofluorescence methods

Dual or triple immunofluorescence procedures were employed to visualize the following markers, cells, and complexes using methods described in our earlier reports (Madhu et al., 2021; Kodali et al., 2021). 1) Proinflammatory microglia positive for IBA-1 and CD68. 2) NLRP3 inflammasome complex positive for IBA-1, NLRP3, and apoptosis-associated speck-like protein containing a CARD (ASC). 3) p62+ structures in the IBA-1+ microglia as an autophagy reporter. 4) phosphorylated S6 ribosomal protein (pS6) expression in the NeuN+ neurons and IBA-1+ microglia to determine the activation of mTOR signaling. Briefly, the sections were washed in phosphate-buffered saline (PBS), blocked with 10% normal donkey serum, and incubated overnight at 4°C in individual primary antibodies (for single immunofluorescence studies) or a cocktail of two or three primary antibodies (for dual and triple immunofluorescence studies). Next, following thorough washing in PBS, the sections were treated with the matching secondary antibodies tagged with fluorescent markers for 60 minutes. Then, the sections were rinsed in PBS and coverslipped with a slow fade/antifade mounting medium (Invitrogen). The primary antibodies employed in the investigation comprised mouse anti-NeuN (1:1000, Millipore Sigma, St. Louis, MO, USA), goat anti-IBA-1 (1:1000, Abcam, Cambridge, MA, USA), rabbit anti-IBA-1 (1:1000, Abcam), mouse anti-CD 68 (1:500, Biorad, Hercules, CA, USA), goat anti-NLRP3 (1:500, Millipore), mouse anti-ASC (1:500, Santa Cruz, Dallas, TX, USA), rabbit anti-phospho-S6 (1:200, cell signaling, Danvers, MA, USA), guinea pig anti-p62 (1:500, Progen, Heidelberg, Germany), rabbit anti-Aβ42 (1:500, Invitrogen), rat anti-dectin-1 (Cleck7a, 1:100, InvivoGen, San Diego, CA, USA). The secondary antibodies used were donkey anti-mouse IgG tagged with Alexa Fluor 488 or 594 (1:200, Invitrogen), donkey anti-goat IgG tagged with Alexa Fluor 488 or 594 (1:200, Invitrogen), donkey anti-rabbit IgG tagged with Alexa Fluor 405 (1:200, Invitrogen), donkey anti-rabbit IgG tagged with Alexa Fluor 488 (1:200, Invitrogen), donkey anti-guinea pig Alexa Fluor 594 (1:200, Jackson ImmunoResearch, West Grove, Pennsylvania, USA), donkey anti-rat IgG tagged with Alexa Fluor 594 (1:200, Invitrogen).

Triple immunofluorescence for NLRP3, ASC, and IBA-1 resulted in detecting the NLRP3 inflammasome complexes in hippocampal and cerebral cortical microglia of naïve, AD, and AD+PLX groups. First, using Z-section analysis in a Leica THUNDER 3D Imager, the total number of NLRP3 inflammasomes (i.e., structures expressing both NLRP3 and ASC) per unit area (∼216 μm^2^) of the CA3 stratum radiatum and cerebral cortex was quantified (3 sections/animal, n=6 animals/group) (Madhu et al., 2021; Attaluri et al., 2022). Next, the percentages of IBA-1+ microglia displaying NLRP3 inflammasome complex (i.e., NLRP3 and ASC co-expression) were measured.

### Quantification of microglia surrounding Aβ plaques

The IBA-1+ microglia surrounding Aβ-42+ plaques in the hippocampus and cerebral cortex were visualized and quantified using 2-μm thick Z-sections from brain tissue sections processed for IBA-1 and Aβ-42 dual immunofluorescence. In both AD and AD-PLX groups, the number of microglia/unit area of the Aβ-42 plaque was computed using three sections per animal (n=4/group). The number of microglia was expressed per 100 μm^2^ of plaque area.

### Visualization of Cleck7a-positive neurodegenerative microglia in and around Aβ plaques

The IBA-1+ microglia expressing Cleck7a (dectin-1, a marker of neurodegenerative microglia [MGnD]) in and around Aβ-42+ plaques were visualized using 1-μm thick Z-sections from brain tissue sections processed for IBA-1, CLEC-7a, and Aβ-42 triple immunofluorescence. In both AD and AD-PLX groups, the extent of Clec7a expression within IBA-1+ microglia in and around Aβ-42 plaques was evaluated (n=6/group).

### Measurement of total microglia and microglia expressing CD68

The number of IBA-1+ microglia in the hippocampus was quantified by stereological counting of IBA1+ cells using every 20^th^ section through the entire hippocampus of naïve, AD, AD+PLX groups (n=6/group). Microglia in the cerebral cortex were also quantified stereologically using every 20^th^ section, but counts were taken as numbers per unit volume (0.1 mm^3^). In addition, we quantified the number of clusters of microglia per unit volume (0.1 mm^3^) of the hippocampus and cerebral cortex in AD and AD+PLX groups (n=6/group) using every 20th section. Stereological counting using StereoInvestigator was performed as described in our previous reports (Hattiangady and Shetty, 2011; Mishra et al., 2015). Percentages of microglia expressing CD68 were quantified from sections processed for IBA-1 and CD68 dual immunofluorescence using Z-section analysis in a confocal microscope as described in our previous study (Kodali et al., 2021). The percentages of IBA-1+ microglia expressing CD68 were computed and compared between naïve, AD, and AD+PLX groups for both hippocampus and cerebral cortex (n=6/group). The percentages of activated microglia were collected from all subfields of the hippocampus (DG, CA1, and CA3) to compile data for the entire hippocampus (3 sections/animal, n=6/group).

### Quantification of astrocyte hypertrophy

The area fractions (AFs) of GFAP+ structures in the hippocampus and cerebral cortex were measured through Image J, using representative sections from all groups (Kodali et al., 2015, 2018). Three sections separated by 600 µm distance were employed in each animal (n=6/group).

### Morphometric analysis of IBA-1+ microglia

Neurolucida (Microbrightfield Inc., Williston, VT) was employed to trace the soma and processes of microglia from the dentate gyrus (DG) of the hippocampus and cerebral cortex, as described in our previous reports (Kodali et al., 2015, 2021a, 2021b). Twenty microglia per DG or cortex (i.e., five microglia/animal, n=6 animals/group) were individually traced in each group using a 100X oil immersion lens. The data, such as total process length, number of nodes, and endings, were computed and compared between naïve, AD, and AD+PLX groups. Sholl’s concentric circle analysis was also employed in the Neurolucida program’s NeuroExplorer component to determine the pattern and extent of processes in microglia from each subregion at various distances from the soma.

### Quantification of NLRP3 inflammasome complex

Triple immunofluorescence for NLRP3, ASC, and IBA-1 detected the NLRP3 inflammasome complexes in hippocampal and cerebral cortical microglia of naïve, AD, and AD+PLX groups. First, using Z-section analysis in a Leica THUNDER 3D Imager, the total number of NLRP3 inflammasomes (i.e., structures expressing both NLRP3 and ASC) per unit area (∼216 μm^2^) of the CA3 stratum radiatum and cerebral cortex was quantified (3 sections/animal, n=6 animals/group) (Madhu et al., 2021; Attaluri et al., 2022). Next, the area fraction (AF) of NLRP3 inflammasome complexes (i.e., structures positive for NLRP3 and ASC) within individual microglia were measured.

### Quantification of mediators and end products of NLRP3 inflammasome activation and additional proinflammatory cytokines

The lysates from the hippocampus and cerebral cortex were utilized for quantifying the mediators and end products of NLRP3 inflammasome activation and additional proinflammatory cytokines using individual enzyme-linked immunosorbent assay (ELISA) kits (n = 6 animals/group: Madhu et al., 2019, 2021). The specific ELISA kits employed in the study include the nuclear subunit of nuclear factor kappa B (NF-kB p65; LSBio, Lynnwood, WA, USA); NLRP3 (Aviva Systems Biology, San Diego, CA, USA), ASC (MyBioSource, San Diego, CA, USA), cleaved caspase-1 (BioVision Inc, Milpitas, CA, USA); interleukin-18 (IL-18), IL-1β (R&D Systems, Minneapolis, MN, USA), tumor necrosis factor-alpha (TNFα; R&D Biosystems), macrophage inflammatory protein-1 alpha (MIP1α; LSBIO), and IL-6 (R&D Biosystems).

### Quantification of ribosomal protein phosphorylated S6 (pS6)

The extent of pS6 expression was quantified within NeuN+ neurons and IBA-1+ microglia in the hippocampus and cerebral cortex using Z-section analysis in a Leica THUNDER 3D Imager (3 images/section, three sections/animal, n=6 animals/group). The pS6 expression in the cytoplasm of neurons and microglia varied within and across groups. Therefore, the AF of pS6+ staining within the soma of neurons and microglia was measured using Image J. The fractions of pS6+ structures in neurons and microglia were measured from all hippocampus subfields (DG, CA1, and CA3) to compile data for the entire hippocampus (3 sections/animal, n=6/group).

### Quantification of phospho-mTOR, pan-mTOR, and ratio of phospho- and pan-mTOR

Tissue lysates from the hippocampus and cerebral cortex were also utilized to quantify the phospho-mTOR and pan-mTOR using an ELISA kit (n = 6 animals/group, Ray Biotech, GA, USA). Next, we determined the ratio of phospho-mTOR and pan-mTOR and compared them across the groups.

### Measurement of p62+ structures

The p62+ structures within microglia and neurons in different hippocampal subfields (the DG, CA1, and CA3 subfields) were quantified. First, the percentages of IBA-1+ microglia expressing p62+ structures in the entire hippocampus were computed and compared between naïve, AD, and AD+PLX groups (3 sections/animal, n=6 animals/group). Next, the area fraction (AF) of p62+ structures within the soma of IBA-1+ microglia and NeuN+ neurons were quantified using Image J. Fourteen microglia from 2-3 images in each animal were measured for each brain region (n=6/group). Neurons were evaluated using ten randomly selected NeuN+ neurons in each cell layer from 2-3 images per animal (n=6/group).

### Quantification of autophagy markers

The lysates from the hippocampus were utilized to quantify the various autophagy markers using ELISA. The kits employed in the study include beclin-1 (Aviva Systems biology, San Diego, CA, USA), autophagy-related gene-5 (ATG-5, MyBioSource), and microtubule-associated protein-1 light chain 3B, (MAP1LC3B, MyBioSource).

### Quantification of Aβ plaques and concentrations of Aβ-42

Aβ plaques in the hippocampus and cerebral cortex were quantified using serial sections (every 15th) and Image J (n=6 animals/group). Serial sections through the posterior half of the hippocampus were employed for the hippocampus since Aβ plaques were minimal in the anterior half of the hippocampus in both AD and AD+PLX groups. Furthermore, we determined the concentrations of Aβ-42 in the hippocampus and cerebral cortex tissue lysates using an ELISA kit from Invitrogen (n=6 animals/group).

### Analysis of hippocampal neurogenesis

The status of hippocampal neurogenesis was measured through the stereological quantification of doublecortin-positive (DCX+) newly born neurons in the subgranular zone-granule cell layer (SGZ-GCL) of the DG using serial sections (every 15^th^, n=6/group) through the entire hippocampus as detailed in our previous reports (Shetty et al., 2020; Rao et al., 2007; Hattiangady et al., 2007).

### Statistical analysis

The animal numbers for immunohistochemical studies per group were determined through a power analysis using G*Power software, using the effect size f as 1.2 (based on our previous data) and alpha as 0.05, which suggested a requirement of final data from a minimum of 4 mice/group to obtain a power of 0.8 and above. However, to obtain robust datasets, we employed n=6/group in all studies. We used one-way ANOVA with Tukey’s post hoc tests to compare data across three groups. We performed the Kruskal Wallis test with Dunn’s post hoc tests when individual groups did not pass the normality test. In comparisons involving two groups, we employed a two-tailed, unpaired Student’s t-test (or a Mann-Whitney U-test when standard deviations differed significantly). A statistically significant value of p<0.05 was employed in all comparisons.

## Results

### Transient CSF1R inhibition in 5xFAD mice depleted microglia by 65% in the hippocampus and cerebral cortex

Immunostaining for IBA-1 visualized the density and distribution of microglia. The examples from the hippocampal and cerebral cortex regions of naïve, AD, and AD+PLX groups are illustrated (Fig. 1 [A-L]). Microglia in AD mice were seen as clusters as well as individual cells (Fig. 1 [B, E, H, K]) in contrast to naïve control mice displaying a homeostatic phenotype, with smaller soma and highly ramified processes (Fig. 1 [A, D, G, J]). Furthermore, microglia within clusters in AD mice displayed the activated phenotype, typified by hypertrophied soma and short processes with no or reduced ramifications (Fig. 1 [E, K]). On the other hand, the residual microglia in AD+PLX mice were mainly scattered and displayed the phenotype of homeostatic microglia, including smaller soma and highly ramified processes (Fig. 1 [C, F, I, L]). Stereological quantification revealed that the total number of microglia significantly increased in the hippocampus and cerebral cortex of the AD group compared to the naïve control group (p<0.01-0.0001; Fig. 1 [M, N]). Moreover, the microglial number was significantly reduced within both hippocampus and cerebral cortex in the AD+PLX group compared to the AD group (65% reduction, p<0.0001) and the naïve control group (p<0.0001, Fig. 1 [M, N]). The reduced number of microglia in the AD+PLX group was also associated with a significant reduction in microglial clusters (40% reduction, p<0.05 – 0.01, Fig. 1 [O, P]).

**Figure 1:**
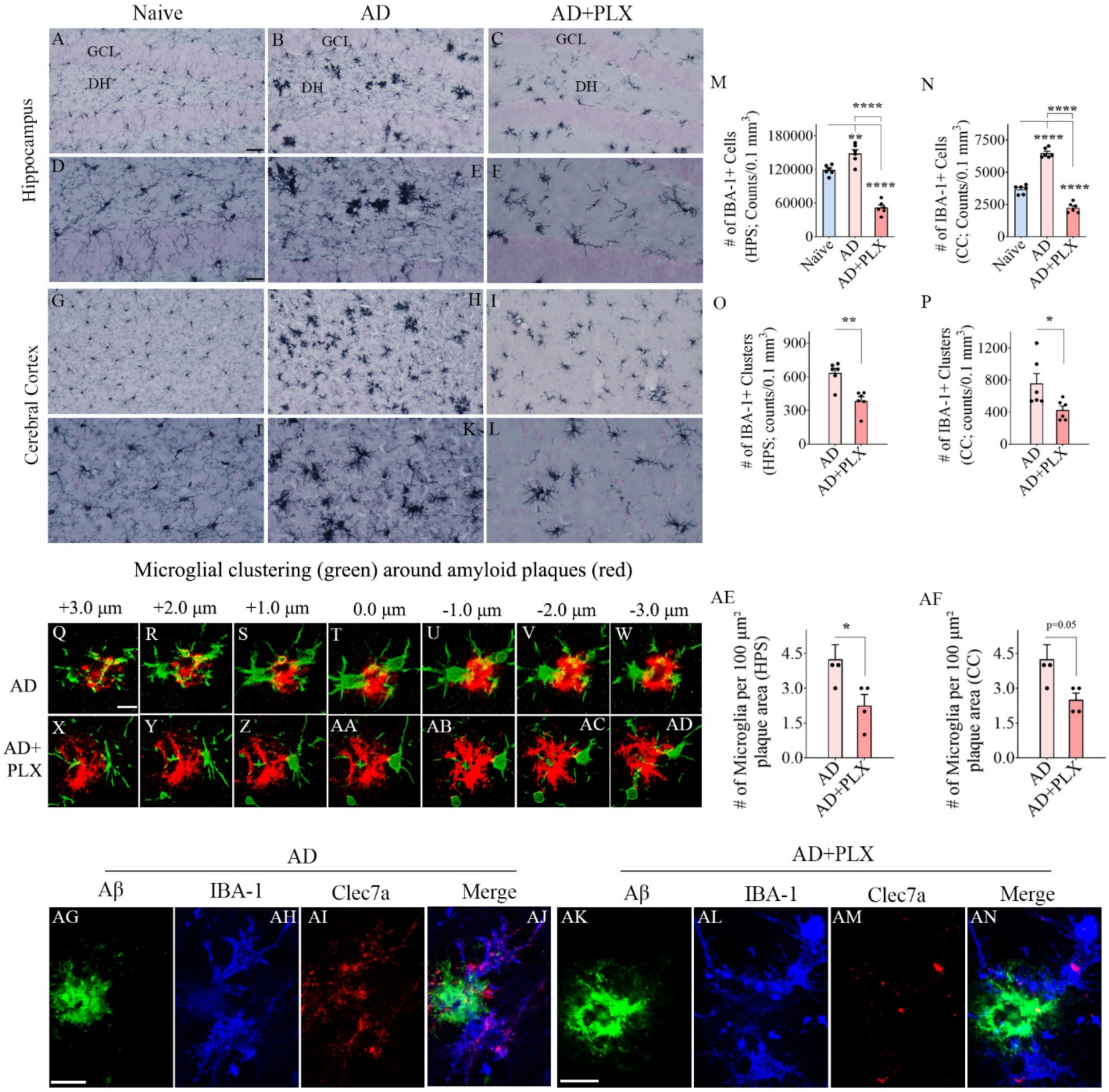
Ten days of CSF1R inhibition in 3-month-old 5xFAD mice resulted in depletion of microglia by 65% in the hippocampus and the cerebral cortex. Figures A–C illustrate examples of IBA1+ microglia from the DG of naïve (A), AD (B), and AD+PLX (C) groups. D-F are magnified views of regions from A-C showing microglial morphology in different groups. Figures G-I illustrate examples of IBA1+ microglia from the cerebral cortex of naïve (G), AD (H), and AD+PLX (I) groups. J-L are magnified views of regions from G-I showing microglial morphology in different groups. The bar charts in M and N compare numbers of IBA-1+ microglia between naive, AD, and AD+PLX groups in the hippocampus (M) and cerebral cortex (N). The bar charts in O and P compare numbers of IBA-1+ microglia clusters between AD and AD+PLX groups in the hippocampus (O) and cerebral cortex (P). Figures Q-AD illustrate confocal images demonstrating microglial clustering around amyloid plaques in the hippocampus from an AD mouse (Q-T) and an AD+PLX mouse (X-AD) groups. The bar charts in AE and AF compare numbers of IBA-1+ microglia per 100 μm^2^ plaque area between AD and AD+PLX groups in the hippocampus (AE) and cerebral cortex (AF). Figures AG-AN illustrate confocal images demonstrating plaque-associated IBA-1+ microglia expressing Clec7a in the hippocampus of an AD mouse (AG-AJ) and an AD-PLX mouse (AK-AN). Scale bar, A-C, G-I = 40 μm; D-F, and J-L = 20 μm; Q-AD; AG-AN = 5 μm. *, p < 0.01; **, p < 0.01; ****, p < 0.0001.

We next investigated plaque-associated microglia using Aβ-42 and IBA-1 dual immunofluorescence in AD and AD+PLX groups (Fig. 1 [Q-AD]). In both groups, microglia were observed around plaques. In the AD group, there was a higher density of cells immediately around the plaques (Fig. 1 [Q-W]) compared to the AD+PLX group (Fig. 1 [X-AD]). Notably, microglia in the AD+PLX group were dispersed around plaques (Fig. 1 [X-AD]). Quantification of microglia per unit area (100 μm^2^) of Aβ-42 plaques revealed a significant reduction in microglial number around plaques in the AD-PLX group vis-à-vis the AD group in the hippocampus (p<0.05, Fig. 1 [AE]) and cerebral cortex (p=0.05, Fig. 1 [AF]). Further evaluation through Aβ-42, IBA-1, and Clec7a triple immunofluorescence revealed varying Clec7a expression within plaque-associated microglia between AD and AD-PLX groups. While all plaque-associated microglia in the AD group displayed robust Clec7a expression [Fig. 1 [AG-AJ]), implying the MGnD phenotype (Krasemann et al., 2017), plaque-associated microglia in the AD-PLX group exhibited greatly reduced Clec7a expression [Fig. 1 [AK-AN]), likely suggesting a non-MGnD phenotype.

### Transient CSF1R inhibition in 5xFAD mice reduced the density of activated microglia in the hippocampus and cerebral cortex

The expression of CD68 within microglia (i.e., activated microglia) was detected through IBA-1 and CD68 dual immunofluorescence and Z-section analysis in a confocal microscope (Supplemental Fig. 1 [A-I, K-S]). Most hypertrophied microglia in AD mice contained large amounts of CD68+ structures, whereas the residual microglia in AD+PLX mice contained reduced CD68+ structures. In the AD group, percentages of IBA-1+ microglia displaying CD68 were higher in the entire hippocampus and the cerebral cortex compared to the naïve control group (p<0.0001, Supplemental Fig. 1 [J, T]). The percentages of CD68+ microglia reduced in the AD+PLX group for the hippocampus and the cerebral cortex (p<0.01-0.0001, Supplemental Fig. 1 [J, T]). Moreover, the percentages of CD68+ microglia in the AD+PLX group were normalized to naïve control levels of the cerebral cortex (p>0.05, Supplemental Fig. 1 [T]). Thus, ten days of PLX treatment in AD mice reduced the extent of activated microglia in both the hippocampus and cerebral cortex.

### Transient CSF1R inhibition in 5xFAD mice transformed the morphology of residual microglia in the hippocampus and cerebral cortex

Tracing of microglia and morphometric analysis using Neurolucida and NeuroExplorer revealed that ten days of PLX treatment in AD mice altered microglial morphology in the hippocampus and cortex (Figs. 2 and Supplemental Fig. 2). Examples of microglia from naïve control, AD, and AD+PLX groups are illustrated for the hippocampal DG (Fig. 2) and cerebral cortex (Supplemental Fig. 2). The naive control group exhibited homeostatic and non-inflammatory phenotypic features (Fig. 2, Supplemental Fig. 2 [A, D]). In contrast, in AD mice, microglia exhibited proinflammatory phenotype (Fig. 2, Supplemental Fig. 2 [B, E]) with reduced process length and a reduced number of nodes and endings (p<0.05-0.0001, Fig. 2, Supplemental Fig. 2 [G-I]). The residual microglia in the AD+PLX group displayed a non-inflammatory phenotype (Fig. 2, Supplemental Fig. 2 [C, F]), akin to that seen in naive control mice, which is evident from increased total process length and a higher number of nodes and endings compared to the AD group (p<0.05-0.0001, Figs. 2, Supplemental Fig. 2 [G-I]). Furthermore, Sholl’s analysis of microglial processes revealed that microglia in the AD group displayed a reduced number of intersections, reduced length of processes, and reduced number of nodes and endings at multiple distances from the soma than the naïve group (Figs. 2, Supplemental Fig. 2 [J-M]). The AD+PLX group exhibited a higher number of intersections, increased length of the processes, and a higher number of nodes and endings at multiple distances from the soma compared to the AD group (p<0.05-0.001, Figs. 2, Supplemental Fig. 2 [J-M]).

**Figure 2:**
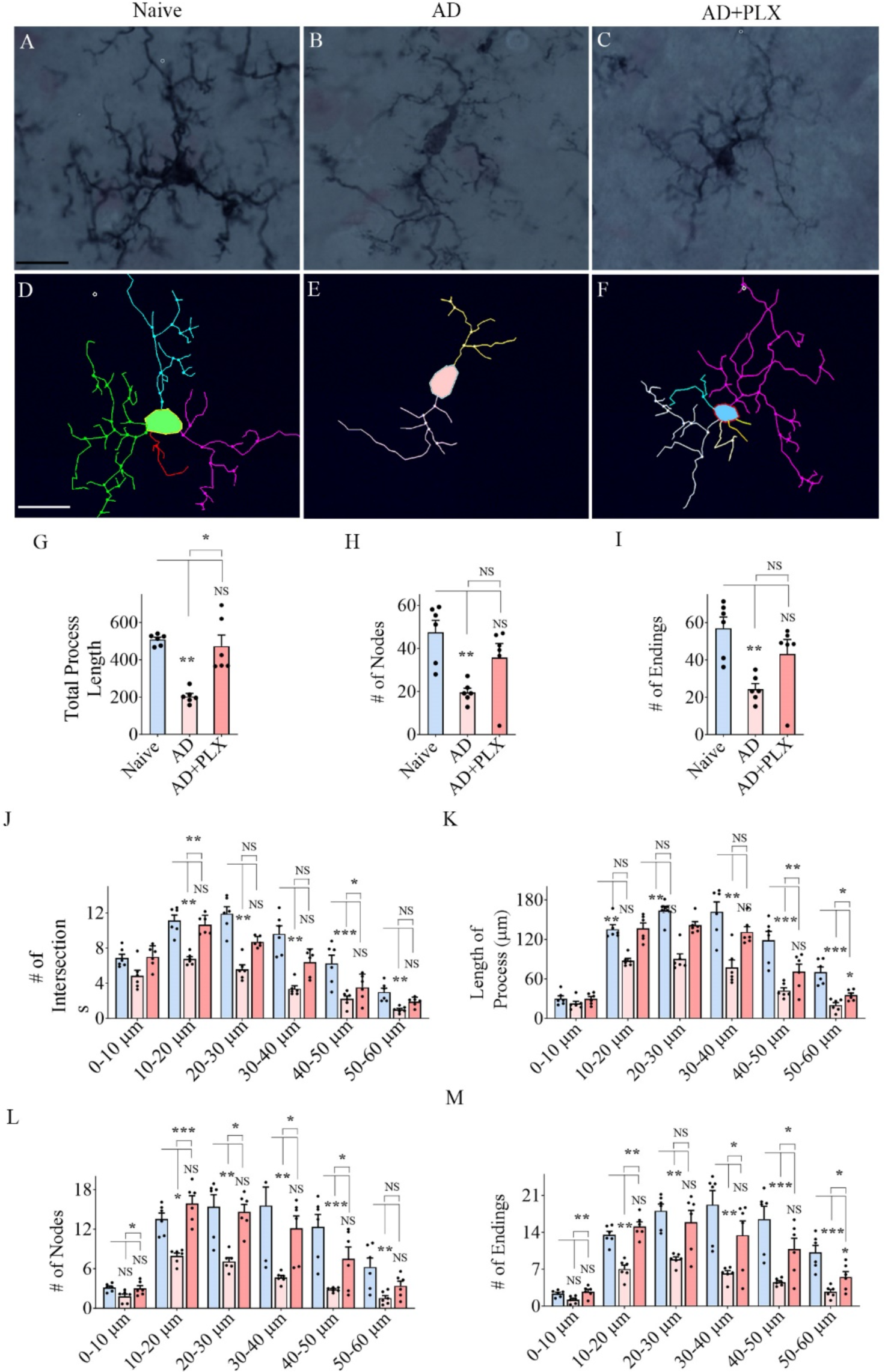
Residual microglia following CSF1R inhibition displayed highly branching and ramified processes in the DG of 5xFAD mice. A-F shows representative examples of microglial morphology traced with Neurolucida from the DG of naïve (A, D), AD (B, E), and AD+PLX (C, F) groups. The bar charts G-I compare the various morphometric measures of microglia between naïve, AD and AD+PLX groups, which include the total process length (G), the number of nodes (H), and the number of process endings (I). The bar charts J-M compare the number of intersections (J), total process length (K), the number of nodes (L), and the number of process endings (M) between naïve, AD and AD+PLX groups at 0–10 μm, 10–20 μm, 20–30 μm, 30–40 μm, 40–50 μm, and 50–60 μm distances from the soma. Scale bar, A-F = 12.5 μm; *, p < 0.05; **, p < 0.01; ***, p < 0.001, NS, not significant.

### Transient CSF1R inhibition in 5xFAD mice reduced NLRP3 inflammasome complexes within residual microglia and diminished NLRP3 inflammasome activation in the hippocampus and cerebral cortex

Z-section analysis of sections processed for IBA-1, NLRP3, and ASC visualized NLRP3 inflammasome complexes (i.e., structures co-expressing NLRP3 and ASC) within IBA-1+ microglia in the hippocampus (Fig. 3 [A-F]) and cerebral cortex (Fig. 4 [A-F]). The presence of NLRP3 inflammasome complex was more frequent in the AD group than in AD+PLX groups (Figs. 3-4 [A-F]). Notably, microglia in the AD-PLX group only occasionally displayed NLRP3 inflammasome complex (Figs. 3-4 [G]). Quantification revealed that the area fraction of inflammasome complexes in microglia was significantly reduced in the AD-PLX group compared to the AD group in the hippocampus and cerebral cortex (p<0.05-0.01, Figs. 3-4 [G]).

**Figure 3:**
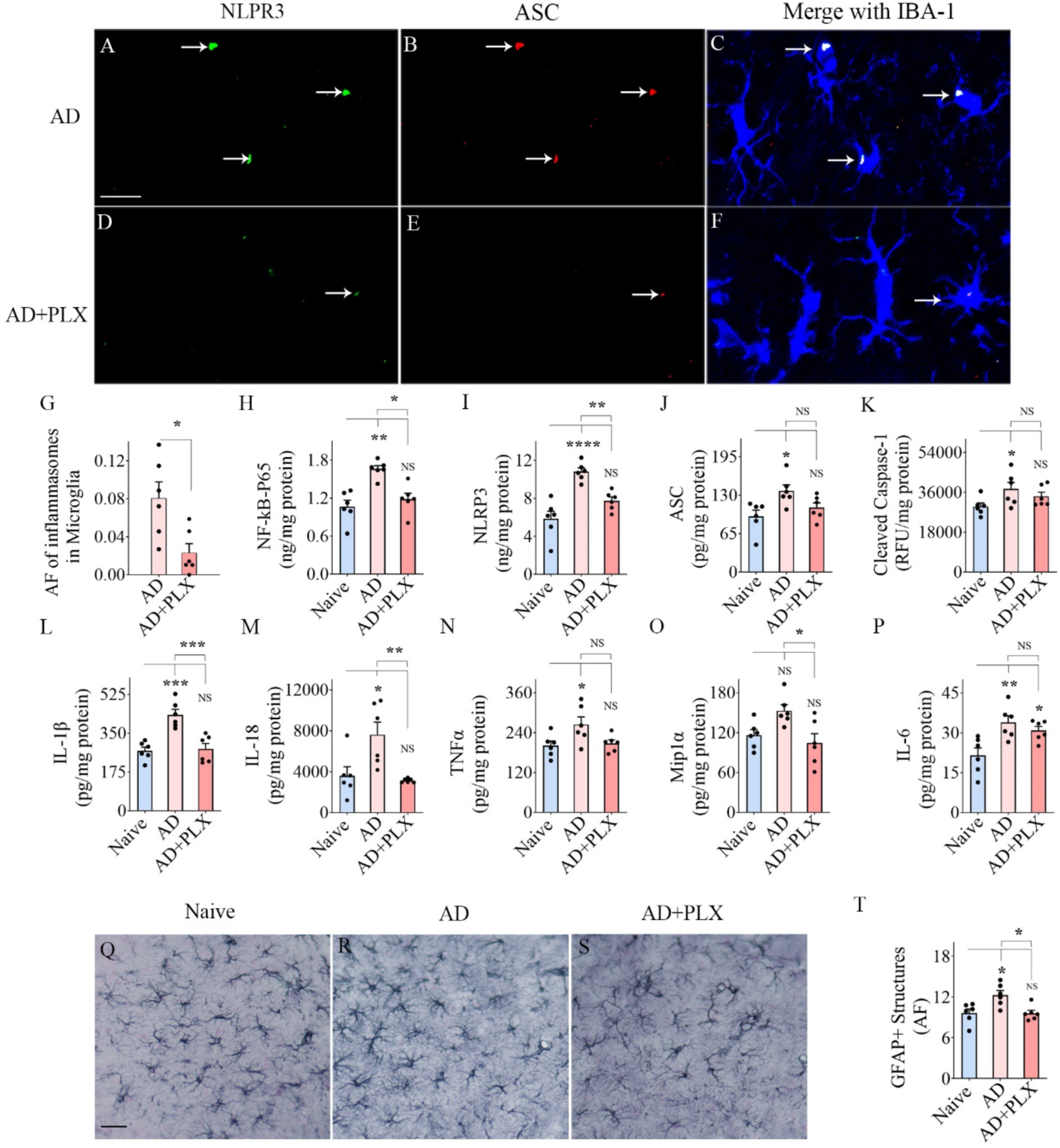
Transient CSF1R inhibition in 5xFAD mice diminished NLRP3 inflammasome complexes in the residual microglia and reduced NLRP3 inflammasome activation, proinflammatory cytokines, and astrocyte hypertrophy in the hippocampus. Figures A–F illustrate nucleotide-binding domain, leucine-rich–containing family, pyrin domain–containing-3 (NLRP3) inflammasome complexes in IBA-1+ microglia from the hippocampus of AD (A-C), and AD+PLX (D-F) groups. The bar chart in G compares the area fraction (AF) of NLRP3 and apoptosis-associated speck-like protein containing a CARD (ASC) complexes in the microglia between AD and AD+PLX groups. Bar chart H compares the concentrations of the nuclear fraction of NF-kB (NF-kB P65) between naïve, AD, and AD+PLX groups. The bar charts I-M compare the concentrations of mediators of NLRP3 inflammasome activation, which include NLRP3 (I), ASC (J), cleaved caspase-1 (K), and end products of NLRP3 inflammasome activation such as interleukin-1 beta (IL-1β; L) and IL-18 (M) between naïve, AD and AD+PLX groups. The bar charts N-P compare the concentrations of other proinflammatory cytokines such as tumor necrosis factor-alpha (TNFα, N), macrophage inflammatory protein (Mip1α; O), and IL-6 (P) between naïve, AD, and AD+PLX groups. Figures Q-S illustrate representative examples of GFAP+ astrocytes in the hippocampus of naïve (Q), AD (R), and AD+PLX (S) groups. Bar chart T compares the AF of GFAP+ structures in the hippocampus between naïve, AD, and AD+PLX groups. Scale bar, A-F = 12.5 μm; Scale bar, Q-S = 40 μm. *, p < 0.05; **, p < 0.01; ***, p < 0.001; ****, p<0.0001; NS, not significant.

**Figure 4:**
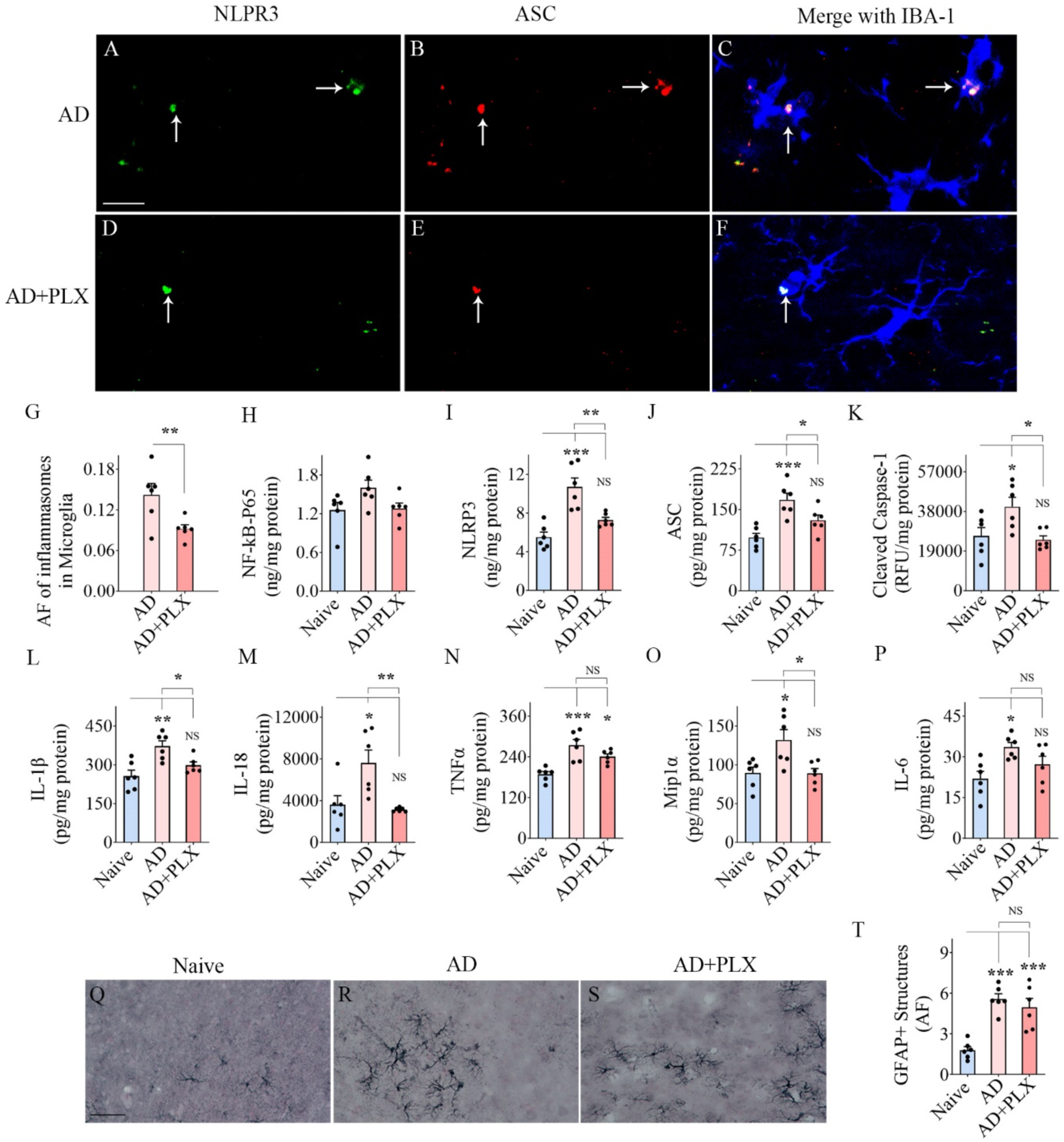
Transient CSF1R inhibition in 5xFAD mice diminished NLRP3 inflammasome complexes in the residual microglia and reduced NLRP3 inflammasome activation and proinflammatory cytokines and astrocyte hypertrophy in the cerebral cortex. Figures A–F illustrate nucleotide-binding domain, leucine-rich–containing family, pyrin domain–containing-3 (NLRP3) inflammasome complexes in IBA-1+ microglia from the cerebral cortex of AD (A-C), and AD+PLX (D-F) groups. The bar chart in G compares the area fraction (AF) of NLRP3 plus apoptosis-associated speck-like protein containing a CARD (ASC) complexes in the microglia between AD, and AD+PLX groups. Bar chart H compares the concentrations of the nuclear fraction of NF-kB (NF-kB P65) between naïve, AD, and AD+PLX groups. The bar charts I-M compare the concentrations of mediators of NLRP3 inflammasome activation, which include NLRP3 (I), ASC (J), cleaved caspase-1 (K), and end products of NLRP3 inflammasome activation such as interleukin-1 beta (IL-1β; L) and IL-18 (M) between naïve, AD and AD+PLX groups. The bar charts N-P compare the concentrations of other proinflammatory cytokines such as tumor necrosis factor-alpha (TNFα, N), macrophage inflammatory protein (Mip1α; O), and IL-6 (P) between naïve, AD, and AD+PLX groups. Figures Q-S illustrate representative examples of GFAP+ astrocytes in the cerebral cortex from the naïve (Q), AD (R), and AD+PLX (S) groups. Bar chart T compares the AF of GFAP+ structures in the cerebral cortex between naïve, AD, and AD+PLX groups. Scale bar, A-F = 12.5 μm; Scale bar, Q-S = 40 μm. *, p < 0.05; **, p < 0.01; ***, p < 0.001; NS, not significant.

Next, we quantified concentrations of proteins implicated in NLRP3 inflammasome activation and proinflammatory cytokines in the hippocampus and the cerebral cortex. AD mice displayed higher levels of proteins that mediate NLRP3 inflammasome activation (NF-kB-p65, NLRP3, ASC, cleaved caspase-1) compared to naïve mice. The differences were significant for all proteins in the hippocampus (p<0.05-0.0001, Fig. 3 [H-K]) and for NLRP3, ASC, and cleaved caspase-1 in the cerebral cortex (p<0.05-0.001, Fig. 4 [I-K]). In contrast, in AD+PLX mice, the concentrations of all these proteins were comparable to naïve control mice in both the hippocampus and cerebral cortex (p>0.05, Figs. 3-4 [H-K]). The concentrations of end products of NLRP3 inflammasome activation (IL-1β and IL-18) and additional proinflammatory cytokines (TNFα, MIP1α, and IL-6) also showed a similar trend. AD mice displayed higher concentrations of these cytokines than naïve mice (p<0.05-0.001, Figs. 3-4 [L-P]), whereas AD+PLX mice exhibited similar concentrations as naïve control mice for all these cytokines in the hippocampus, and IL-1β, IL-18, MIP1α, and IL-6 in the cerebral cortex (p>0.05, Figs. 3-4 [L-M, O-P]). Ten days of PLX treatment in AD mice diminished the concentration of various proteins involved in NLRP3 inflammasome activation and multiple downstream proinflammatory cytokines.

### Transient CSF1R inhibition in 5xFAD mice reduced astrocyte hypertrophy in the hippocampus

Examples of GFAP immunoreactive astrocytes in the CA3 subfield and cerebral cortex (Figs. 3-4 [Q-S]) are illustrated. Quantification using Image J revealed that AF of astrocyte elements in the hippocampal CA3 region and cerebral cortex of the AD group were elevated compared to the naïve group (p<0.05-0.001; Figs. 3-4 [T]). In the AD+PLX group, the AF of astrocyte elements was comparable to the naïve group in the hippocampus (Fig. 3 [T]) but not in the cerebral cortex (Fig. 4 [T]), where astrocyte elements remained comparable to the AD group (p>0.05). Thus, ten days of PLX treatment in AD mice reduced astrocyte hypertrophy in the hippocampus but not in the cerebral cortex.

### Transient CSF1R inhibition in 5xFAD mice diminished mTOR signaling in the hippocampus and cerebral cortex

mTOR signaling within hippocampal and cerebral cortical neurons and microglia were visualized and quantified through NeuN/pS6 and IBA-1/pS6 dual immunofluorescence, Z-section and Image J analyses (Figs. 5-6 [A-F, H-M]). pS6 is one of the primary downstream targets and the effector of the mTOR pathway and hence serves as a marker for mTOR activity (Leontieva et al., 2012; Morgan-Warren et al., 2013). AF analysis of individual neurons revealed diminished pS6 expression within hippocampal and cerebral cortical neurons in the AD+PLX group compared to the AD group (Figs. 5-6 [G]). Such reduction was statistically significant in the cerebral cortex (p<0.01, Fig. 6 [G]), implying reduced mTOR signaling. AF analysis of pS6+ immunoreactivity within individual microglia also revealed decreased pS6 expression in the AD+PLX group compared to the AD group in the hippocampus and cerebral cortex (p<0.05-0.01, Fig. 5-6 [N]). Thus, microglia depletion reduced mTOR signaling within both neurons and microglia. To further probe the mTOR signaling alteration, we measured the concentrations of pan-mTOR, phospho-mTOR, and the ratio of phospho- and pan-mTOR in the hippocampus and the cerebral cortex (Figs. 5-6 [O-Q]). The ratio of phospho- and pan-mTOR levels was upregulated in the AD group compared to the naive group in both the hippocampus and the cerebral cortex (p<0.01, Figs. 5-6 [Q]), but the AD+PLX group displayed a similar ratio of phospho- and pan-mTOR as the naive group (p>0.05, Figs. 5-6 [Q]), implying reduced mTOR signaling in both regions. Thus, ten days of PLX treatment in AD mice can significantly dampen mTOR signaling in both the hippocampus and cerebral cortex.

**Figure 5:**
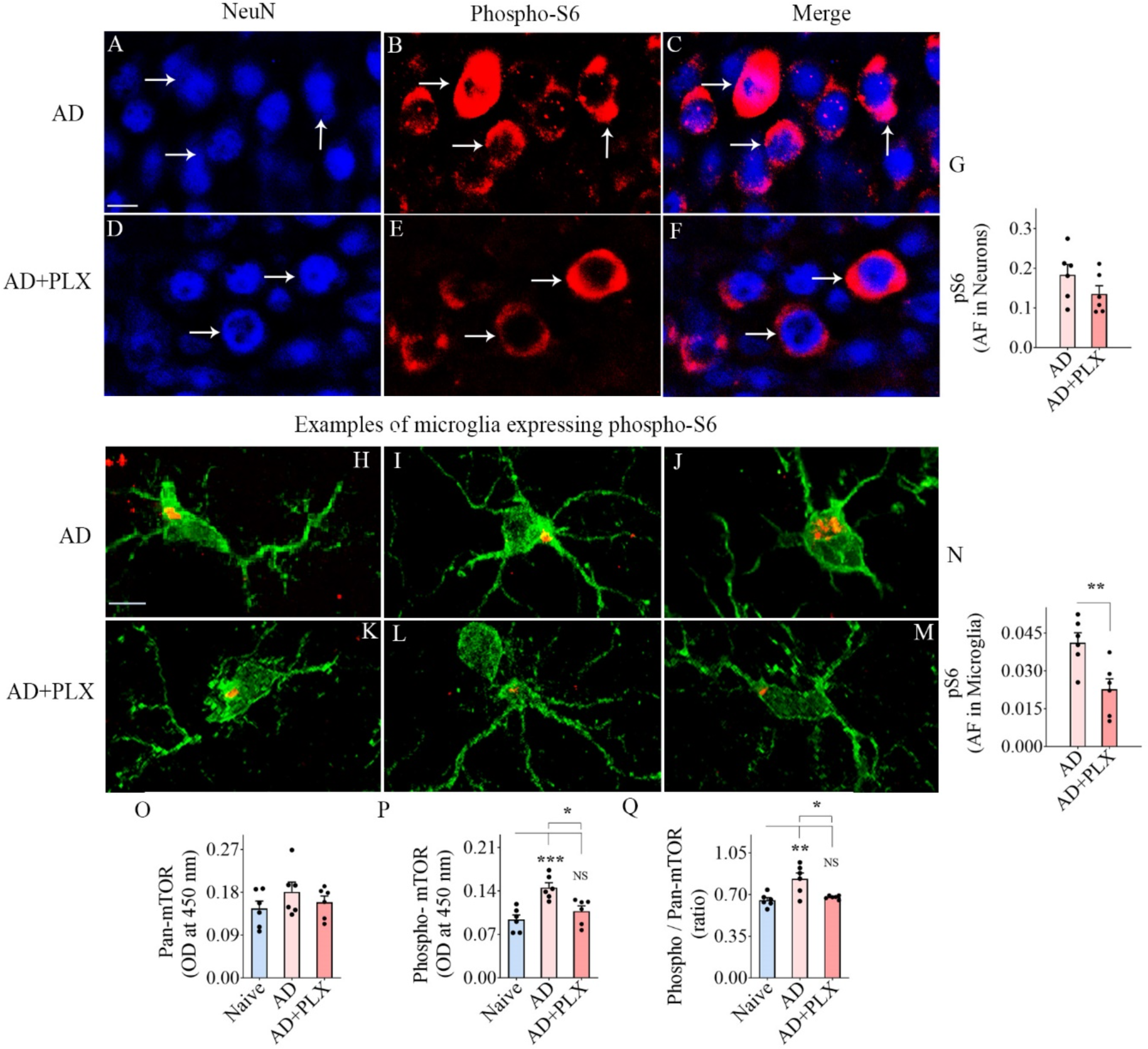
Ten days of CSF1R inhibition in 5xFAD mice reduced phospho-S6 (pS6, a measure of mechanistic target of rapamycin [mTOR]) in hippocampal neurons and microglia and the entire hippocampus. Figures A-F illustrate hippocampus CA3 pyramidal neurons expressing pS6 from AD (A-C) and AD+PLX (D-F) groups. The bar chart G compares the area fraction (AF) of pS6 in NeuN+ CA3 pyramidal neurons between AD and AD+PLX groups. Figures H-M illustrate hippocampal microglia expressing pS6 from AD (H-J) and AD+PLX groups (K-M) groups. The bar chart N compares the AF of pS6 in IBA-1+ hippocampal microglia between AD and AD+PLX groups. The bar charts O-Q compare the concentrations of pan-mTOR (O), phospho-mTOR (P), and the ratio of phospho/pan-mTOR (Q) between naïve, AD and AD+PLX groups in the hippocampus. Scale bar, A-F=10 μm, H-M = 2.5 μm.*, p<0.05; **, p<0.01; ***, p < 0.001; NS, not significant.

**Figure 6:**
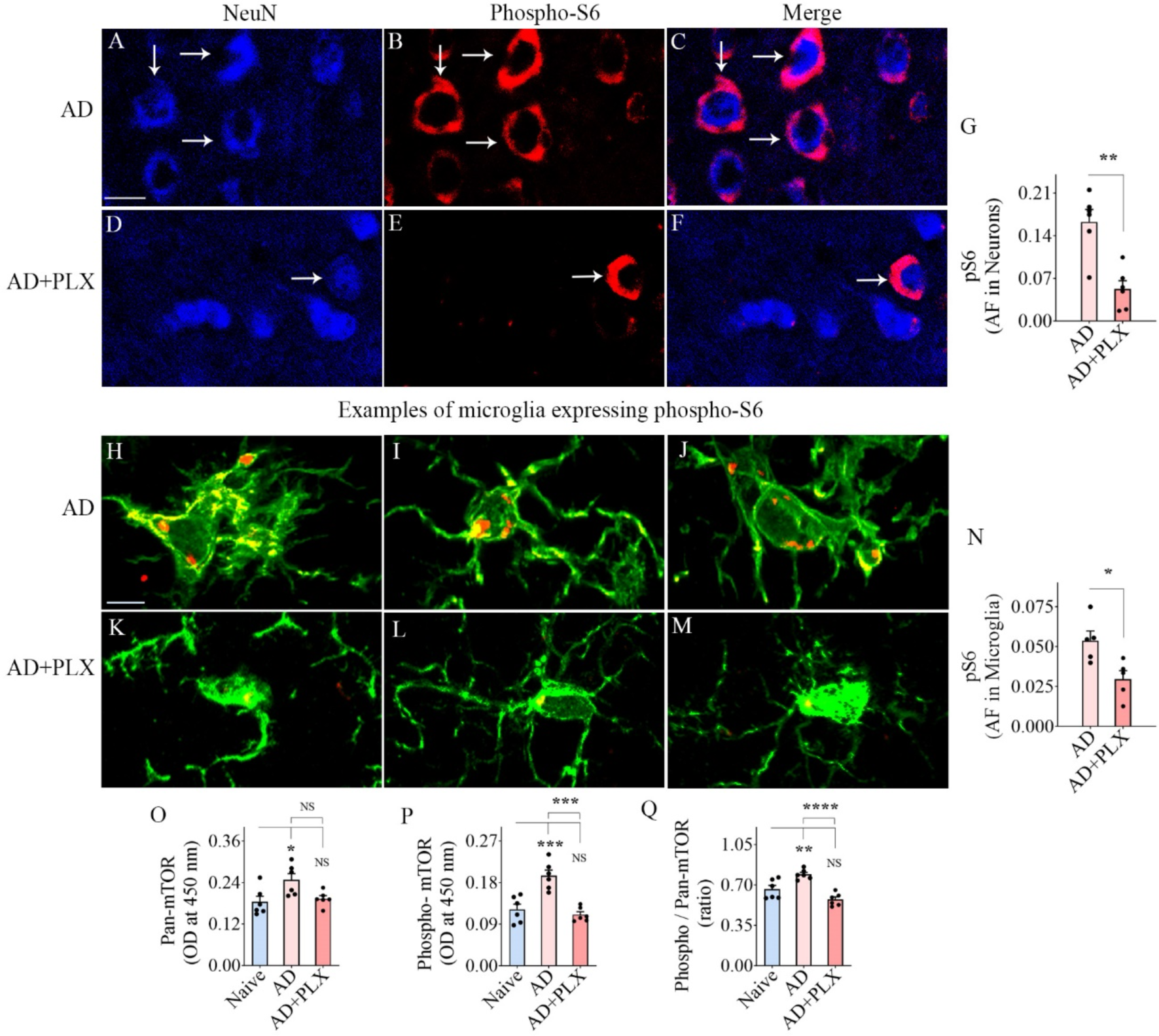
Ten days of CSF1R inhibition in 5xFAD mice reduced phospho-S6 (pS6, a measure of mechanistic target of rapamycin [mTOR]) in cerebral cortical neurons and microglia and the cerebral cortex. Figures A-F illustrate neurons from the cerebral cortex expressing pS6 from AD (A-C), and AD+PLX groups (D-F) groups. The bar chart G compares the area fraction (AF) of pS6 in NeuN+ neurons between AD and AD+PLX groups. Figures H-M illustrate microglia expressing pS6 from AD (H-J) and AD+PLX groups (K-M) groups. The bar chart N compares the AF of pS6 in IBA1+ microglia between AD and AD+PLX groups. The bar charts O-P compare the concentrations of pan-mTOR (O), phospho-mTOR (P), and the ratio of phospho/pan-mTOR (Q) between naïve, AD and AD+PLX groups in the cerebral cortex. Scale bar, A-F=10 μm, H-M = 2.5 μm. *, p<0.05; **, p<0.01; ***, p < 0.001; ****, p < 0.0001; NS, not significant.

### Transient CSF1R inhibition in 5xFAD mice enhanced autophagy in the hippocampus and cerebral cortex

Dual immunofluorescence for IBA-1/p62 and NeuN/p62 and Z-section analysis and quantification via Image J revealed the extent of autophagy within neurons (Figs. 7-8 [A-F]) and microglia (Fig. 7-8 [H-M]) in the hippocampus and cerebral cortex. The p62 protein serves as a classical marker of autophagic flux because it accumulates with autophagic cargo but declines when autophagy is active (Pankiv et al.,2007; Liu et al., 2019). Compared to the AD group, the AD-PLX group displayed reduced AFs of p62 within hippocampal and cerebral cortex neurons (Figs. 7-8 [G]). Such reductions were statistically significant in the cerebral cortex (p<0.01, Fig. 8 [G]). Analysis of the p62 expression in microglia also revealed a similar trend. The AD+PLX group exhibited reduced p62 expression compared to the AD group in the hippocampus and cerebral cortex (p>0.001-0.0001, Fig. 7-8 [N]). Overall, p62 expression in neurons and microglia implied improved autophagy in the AD+PLX group. Consistent with these results, we found decreased concentrations of beclin-1 and ATG-5 in the hippocampus and cerebral cortex of the AD group compared to the naive group (Figs. 7-8 [O]), and the decrease was significant for beclin-1 in the hippocampus (Figs. 7 [O]). Moreover, both beclin-1 and ATG-5 concentrations in the AD+PLX group were comparable to the naive group in the hippocampus (p>0.05, Fig. 7 [O-P]) and significantly higher than the AD group (p<0.01, Fig. 7 [O-P]). Measurements from the cerebral cortex showed a similar trend, but the differences were not statistically significant. Thus, partial microglia depletion normalized autophagy within the leftover microglia in the hippocampus of 5xFAD mice to naive control levels.

**Figure 7:**
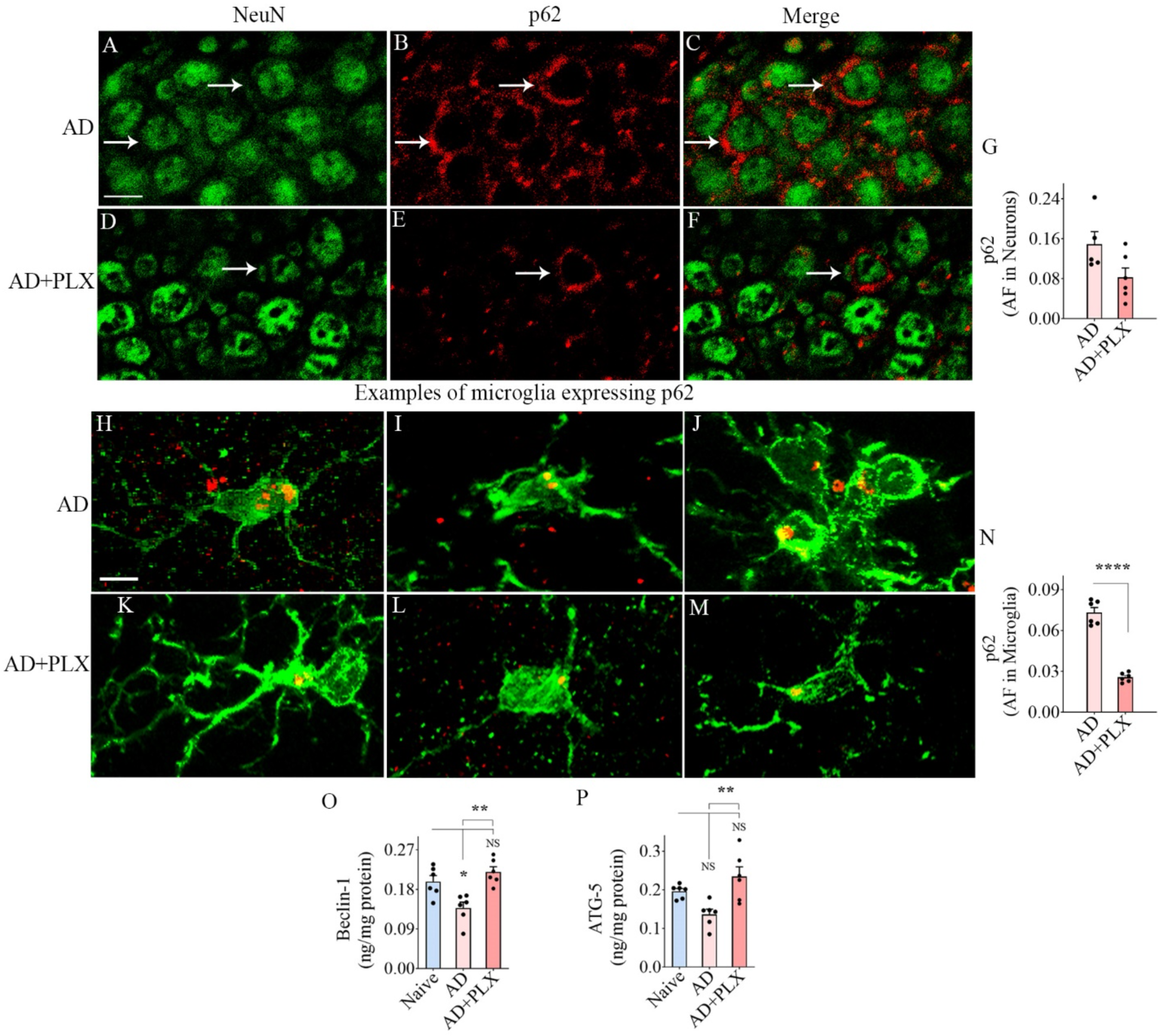
Ten days of CSF1R inhibition in 5xFAD mice enhanced autophagy in the hippocampal neurons, microglia and the hippocampus. Figures A-F show examples of neurons expressing p62 (a marker of autophagic flux) in the CA3 subregion of the hippocampus of AD (A-C) and AD+PLX (D-F) groups. The bar chart G compares the area fraction (AF) of pS6 in NeuN+ neurons between AD and AD+PLX groups. Figures H-M illustrate microglia expressing pS6 from AD (H-J) and AD+PLX groups (K-M) groups. The bar chart N compares the AF of pS6 in IBA-1+ microglia between AD and AD+PLX groups in the hippocampus. The bar charts O-P compare the concentration of autophagy-related proteins, beclin-1 (O), and autophagy-related 5 (ATG-5, P), between naïve, AD and AD+PLX groups. Scale bar, A-F = 10 μm, H-M = 2.5 μm *, p<0.05; **, p<0.01; ****, p < 0.0001; NS, not significant.

**Figure 8:**
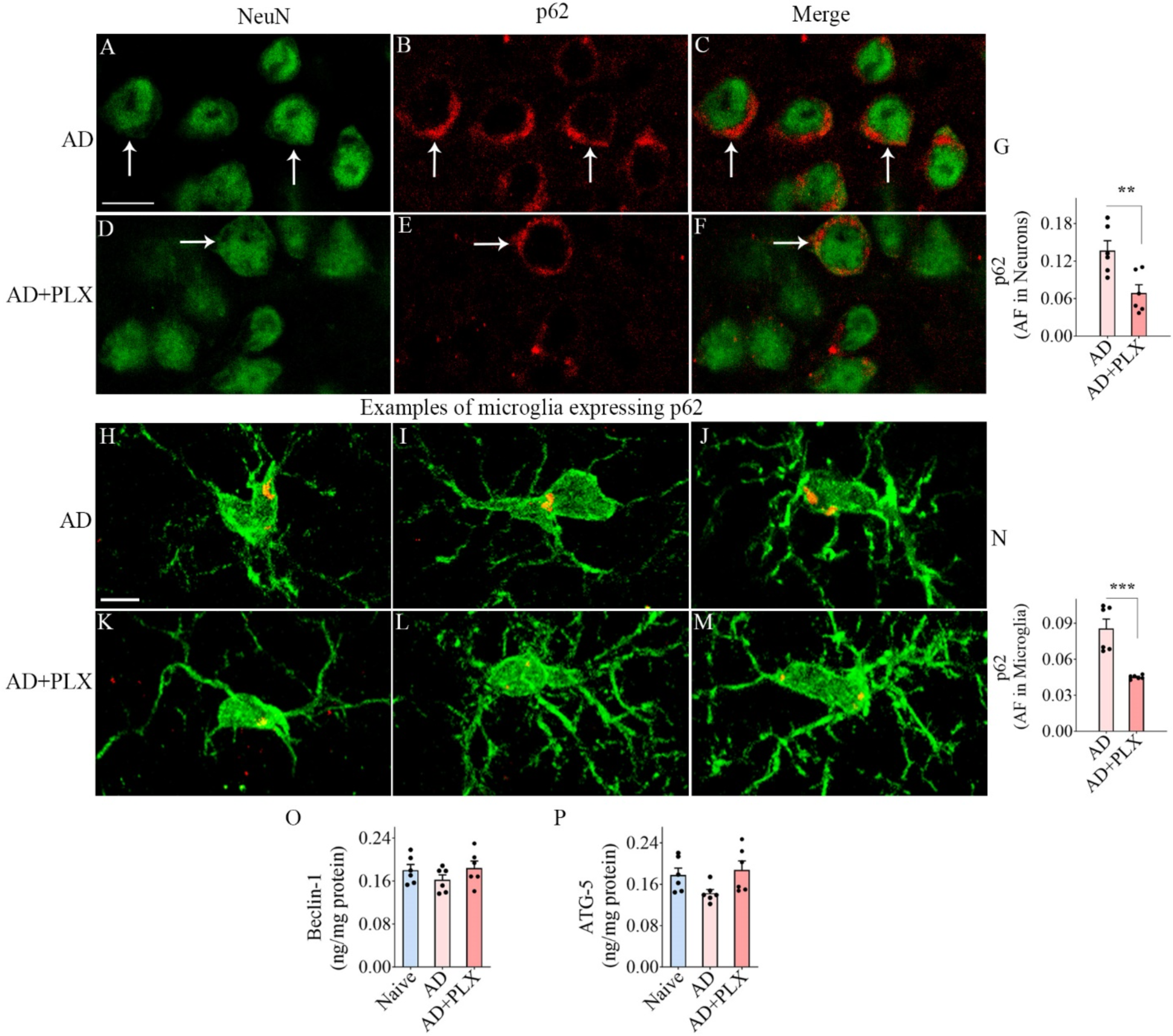
Ten days of CSF1R inhibition in 5xFAD mice enhanced autophagy in the cortical neurons, microglia, and the cortex. Figures A-F show examples of neurons expressing p62 in the cerebral cortex of AD (A-C) and AD+PLX (D-F) groups. The bar chart G compares the area fraction (AF) of pS6 in NeuN+ neurons between AD and AD+PLX groups. Figures H-M illustrate microglia expressing pS6 from AD (H-J) and AD+PLX groups (K-M) groups. The bar chart N compares the AF of pS6 in IBA-1+ microglia between AD and AD+PLX groups. The bar charts O-P compare the concentration of autophagy-related proteins, beclin-1 (O), and autophagy-related 5 (ATG-5, P), between naïve, AD and AD+PLX groups in the cortex. Scale bar, A-F = 10 μm, H-M = 2.5 μm. **, p<0.01; ***, p < 0.001; NS, not significant.

### Transient CSF1R inhibition in 5xFAD mice did not alter Aβ plaques but reduced Aβ-42 concentration

Examples of Aβ plaques from the hippocampus and cerebral cortex are illustrated (Fig. 9 [A-D]). In these regions, the density of Aβ plaques appeared similar between AD and AD+PLX groups. Quantification of AFs of Aβ plaques in these brain regions using Image J confirmed that the extent of Aβ plaques in the AD group matched the AD+PLX group (p>0.05, Fig. 9 [E-F]). Furthermore, the concentrations of soluble Aβ-42 were also not statistically significant between the AD and AD+PLX groups in both the hippocampus and cerebral cortex (p>0.05, Fig. 9 [G-H]). Thus, ten days of PLX treatment in AD mice did not significantly reduce the extent of Aβ plaques and the soluble Aβ-42 in the hippocampus and cerebral cortex.

**Figure 9:**
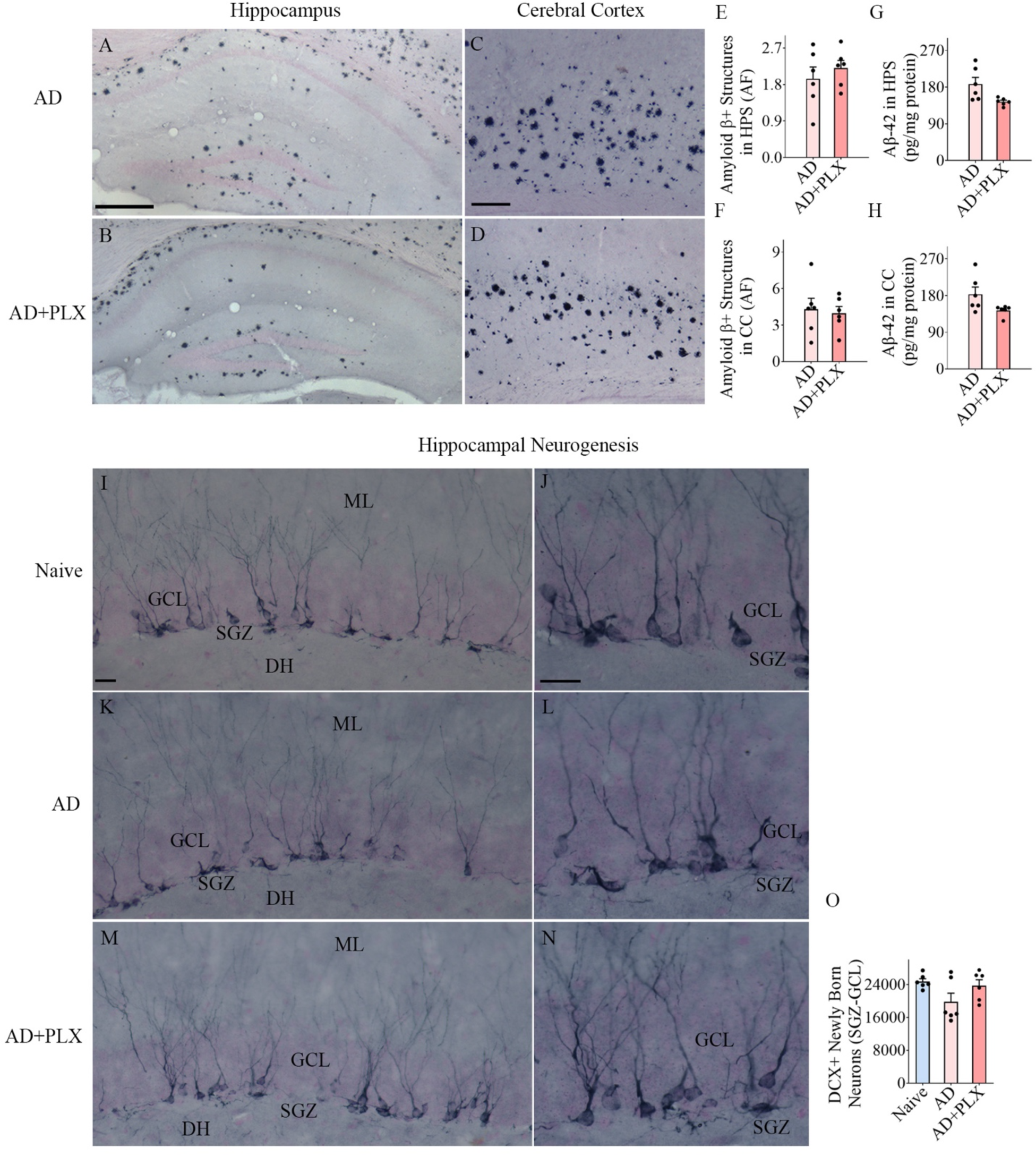
Ten days of CSF1R inhibition in 5xFAD mice did not alter the amyloid-beta plaques in the hippocampus and cerebral cortex and hippocampal neurogenesis. Figures A–D illustrate examples of amyloid beta (Aβ) plaques in the hippocampus (A, B) and the cerebral cortex (C, D) from AD (A, C) and AD+PLX (B, D) groups. The bar charts E-F compare the area fraction of Aβ plaques between AD and AD+PLX groups from the hippocampus (E) and the cerebral cortex (F). Scale bar, A-B = 250 μm, C-D=125 μm. Bar charts G-H compare Aβ-42 concentrations between AD and AD+PLX groups in the hippocampus (G) and the cerebral cortex (H). Figures I-N illustrate examples of doublecortin-positive (DCX+) newly born neurons from naïve (I-J), AD (K-L), and AD+PLX (M-N) groups. Bar chart O compares the number of DCX+ neurons between naïve, AD, and AD+PLX groups. GCL, granule cell layer; SGZ, subgranular zone; DH, dentate hilus; ML, molecular layer. Scale bar, L-N = 20 μm.

### Transient CSF1R inhibition in 5xFAD mice did not impact hippocampal neurogenesis

The extent of hippocampal neurogenesis was ascertained through DCX immunostaining (Fig. 9 [I-N]). Stereological quantification of DCX+ newly born neurons in the SGZ-GCL revealed no differences between naïve, AD, and AD+PLX groups (p>0.05, Fig. 9 [O]). Thus, ten days of PLX treatment in AD mice did not alter hippocampal neurogenesis.

## Discussion

The study provides novel evidence that 10-day CSF1R inhibition in three-month-old 5xFAD mice, representing the early stage of neuroinflammation in AD, results in ∼65% depletion of microglia in the hippocampus and cerebral cortex, with the residual microglia predominantly presenting a non-inflammatory phenotype. The non-inflammatory phenotype of leftover microglia was evident from highly branched and ramified processes with reduced NLRP3 inflammasome complexes, mTOR signaling, and enhanced autophagy. Moreover, the altered microglial phenotype following 10-day CSF1R inhibition was found to be associated with decreased mTOR signaling, increased autophagy in neurons, and reduced astrocyte hypertrophy while showing no significant changes in the extent of Aβ-42 plaques, soluble Aβ-42, or hippocampal neurogenesis.

### Potential reasons for leftover microglia displaying a predominantly noninflammatory phenotype

A series of assessments has verified that the residual microglia in the hippocampus and cerebral cortex of 5xFAD mice following ten days of CSF1R inhibition predominantly displayed a noninflammatory phenotype. These assessments encompassed morphometric analyses quantifying the intricacies of microglial processes, such as branches, nodes, and process lengths at various distances from the soma, illustrating the resemblance of residual microglial processes to those of the naive control group. Additional observations supporting the predominant elimination of activated microglia in AD mice receiving ten days of CSF1R inhibition include the AD group receiving a standard diet displaying large numbers of microglial clusters comprising microglia with hypertrophied soma, thicker and shorter processes, and increased incidence of NLRP3 inflammasome complexes. In contrast, in the AD+PLX group, such clusters were reduced, and the dispersed residual microglia displayed diminished NLRP3 inflammasome complexes. Moreover, while microglia were found around plaques in both AD and AD+PLX groups, there were significantly fewer plaque-associated microglia in the AD+PLX group compared to the AD group. Additional evaluation through Aβ-42, IBA-1, and Clec7a triple immunofluorescence demonstrated that all plaque-associated microglia in the AD group exhibited pronounced expression of Clec7a, indicative of an MGnD phenotype (Krasemann et al., 2017). Conversely, microglia associated with plaques in the AD-PLX group displayed diminished Clec7a expression, suggesting a non-MGnD phenotype. There was also an overall reduction in proinflammatory milieu in the hippocampus and cerebral cortex, which could be gleaned from reduced concentrations of NLRP3 inflammasome activation mediators (NF-kB-p65, NLRP3, ASC, and cleaved caspase-1), end products (IL-1β and IL-18), and other proinflammatory cytokines such as TNFα, MIP1α, and IL-6 in the hippocampus and cerebral cortex of AD-PLX group compared to the AD group. These results are consistent with the diminished proinflammatory cytokine concentrations seen after CSF1R inhibition in another model of AD (Mancuso et al., 2019).

The exact reasons for leftover microglia displaying a predominantly noninflammatory phenotype in PLX-treated 5xFAD mice are unclear. Short-term CSF1R inhibition in the early stage of AD predominantly ablates activated microglia, possibly due to a higher demand for CSF1 for survival by such microglia. Although direct evidence supporting enhanced CSF1R expression in activated microglia due to a higher demand for CSF1 is lacking, an overall CSF1R upregulation has been observed in mouse models of amyloidosis and neurotoxic models of Parkinson’s disease (Murphy et al., 2000; Neal et al., 2020) and post-mortem samples from AD and Parkinson’s disease (PD) patients (Akiyama et al., 1994; Neal et al., 2020). Also, a study has shown that microglial proliferation increases progressively near Aβ plaques in the APP/PS1 AD model (Olmos-Alonso et al., 2016), which may be mediated through increased CSF1R in plaque-associated microglia. Repolarizing the activated microglia into a homeostatic phenotype after CSF1R inhibition has also been observed in models of multiple sclerosis (Nissen et al., 2018) and Parkinson’s disease (Neal et al., 2020). Regardless of the reasons, the data support the increased vulnerability of activated microglia to CSF1R inhibition, at least in the early stage of AD. It remains to be investigated whether a similar elimination of activated microglia would occur in the later stages of AD. In this context, a study in ∼5.5 months old tau-seeded 5xFAD mice showing that 45 days of CSF1R inhibition preferentially eliminates non-plaque-associated microglia vis-à-vis plaque-associated microglia is relevant (Lodder et al., 2021). The discrepancy between the latter study and the current study regarding the type of microglia eliminated following CSF1R inhibition likely reflects the age of the mice, disease stage, the extent of microglial activation, and the duration of CSF1R inhibition.

### Implications of reduced NLRP3 inflammasome activation in residual microglia after 10-day CSF1R Inhibition

Reductions in NLRP3 inflammasome complexes in the residual microglia following ten days of CSF1R inhibition in the early stage of AD are beneficial because NLRP3 inflammasome activation perpetuates chronic neuroinflammation and progression of AD pathogenesis (Ising et al., 2019; Bai et al., 2021). The NLRP3 inflammasome complex comprises NLRP3, ASC, and caspase-1 precursor proteins. Increases in NLRP3, a NOD-like receptor protein with pattern recognition, recruit caspase-1 precursor protein by binding to ASC through the pyrin domain at the N-terminal, which activates caspase-1. Activation of caspase-1 facilitates the maturation of pro-IL-1β and pro-IL-18 into mature IL-1β and IL-18. The secreted mature IL-1β and IL-18 contribute to neurodegeneration through downstream inflammatory cascades (Franchi et al., 2009; Zhang et al., 2017). The events that upregulate NLRP3 in AD include increased levels of the danger-associated molecular patterns (DAMPs) such as adenosine triphosphate, reactive oxygen species, and cathepsin B (Savage et al., 2012), which activate nuclear factor kappa B through toll-like receptors (Yang et al., 2020). Mitochondrial DNA released from the damaged mitochondria can also directly upregulate NLRP3 in AD via potassium outflow or calcium influx (Wyss-Coray, 2006; Murphy et al., 2012). In addition, activation of NLRP3 inflammasomes in AD can occur by fibrillar Aβ aggregates, Aβ oligomers, and protofibrils (Friker et al., 2020).

While activation of NLRP3 inflammasome is beneficial to the organism in certain conditions as it can restrain microbial infection or endogenous cell damage, its overactivation is detrimental in AD (Hanslik and Ulland, 2020; Ising et al., 2019; Heneka et al., 2019). Overactivation of the NLRP3 inflammasomes in microglia transforms them into a proinflammatory phenotype, resulting in reduced phagocytosis of Aβ-42 by microglia, which can lead to enhanced Aβ deposition and progression of AD pathogenesis (Lučiūnaitė et al., 2020). Such effects are also apparent in microglia with reduced NLRP3 inflammasome activation, displaying a noninflammatory phenotype and exhibiting increased phagocytosis of Aβ (Cherry et al., 2014). Moreover, NLRP3 inflammasome overactivation promotes the pathological formation of tau protein, which can accelerate AD-related neurodegeneration (Ising et al., 2019). Therefore, many studies have focused on inhibiting NLRP3 inflammasome activation in AD to suppress chronic neuroinflammation and improve brain function (Bai et al., 2021; Feng et al., 2020; Lonnemann et al., 2020; Milner et al., 2021; Van Zeller et al., 2021). For example, reduction of caspase-1 and IL-1β through knockdown of NLRP3 in AD models can considerably eliminate Aβ-42 (Wu et al., 2017), and functional loss of the NLRP3 inflammasome could diminish hyperphosphorylation and aggregation of tau by regulating tau kinase and phosphorylase (Dempsey et al., 2017). Thus, short-term CSF1R inhibition resulting in reduced NLRP3 inflammasome complexes in residual microglia in the early stage of AD, along with an overall reduction in concentrations of NLRP3 inflammasome activation mediators and end products observed in this study, promises to maintain better brain function for extended periods. However, it remains to be addressed whether short-term CSF1R inhibition at regular intervals (e.g., ten days of CSF1R inhibition every 6-8 weeks) would postpone cognitive and memory problems of AD for extended periods.

### Significance of reduced mTOR signaling and enhanced autophagy in residual microglia after 10-day CSFIR Inhibition

A predominant elimination of activated microglia in AD mice receiving ten days of CSF1R inhibition led to decreased mTOR signaling in neurons in the cerebral cortex and microglia in both the hippocampus and the cerebral cortex. Moreover, the overall mTOR signaling, indicated by concentrations of p-mTOR and the ratio of p-mTOR and pan-mTOR, decreased in AD mice receiving ten days of CSF1R inhibition. mTOR hyperactivation has been seen in both animal models of AD and post-mortem brain samples from AD patients (Li et al., 2005; Caccamo et al., 2010, 2014). Increased mTOR signaling in AD increases Aβ production and promotes aggregation by directly inhibiting autophagy (Zhao et al., 2015; Caccamo et al., 2011). The precise reason for decreased mTOR signaling with the predominant elimination of activated microglia in the early stage of AD observed in this study is unknown. It could be due to an overall decrease in NLRP3, as studies have shown that NLRP3 is a binding partner of mTOR, and NLRP3 silencing reduces the phosphorylation of mTOR (Cosin-Roger et al., 2017). Thus, reduced NLRP3 levels, as observed in this study, diminished the physical interaction between NLRP3 and mTOR with a net effect of diminished mTOR signaling. One of the benefits of decreased mTOR signaling is the resulting increase in autophagy. Indeed, increased autophagy in the AD-PLX group in this study was apparent in hippocampal microglia and cerebral cortical neurons and microglia from diminished p62 expression. Moreover, compared to the AD group, there was an overall increase in autophagy in the hippocampus of the AD-PLX group, indicated by higher levels of autophagy enhancers like Beclin-1 and ATG-5. In this study, the increased autophagy in the AD-PLX group is likely due not only to reduced mTOR signaling but also to lower levels of proinflammatory cytokines in the environment, as elevated proinflammatory cytokines like TNF-α can impede autophagic flux in microglia (Jin et al., 2018). Thus, the reduced NLRP3 inflammasome activation, decreased mTOR signaling, and increased autophagy in residual microglia after 10-day CSF1R inhibition are closely connected.

Enhanced autophagy in residual microglia is beneficial because it helps remove NLRP3 inflammasome activators, such as intracellular DAMPs, NLRP3 inflammasome components, and cytokines (Ma et al., 2013; Galluzzi et al., 2017). Conversely, dysfunctional autophagy can result in inflammatory diseases due to excessive inflammasome activation (Levine et al., 2011; Deretic et al., 2013). Therefore, maintaining increased autophagy in AD is crucial for reducing the hyperinflammatory response. Enhanced autophagy in microglia may also improve the phagocytosis of Aβ-42 (Plaza-Zabala et al., 2017; Lucin et al., 2013). However, the current study did not find reductions in Aβ plaques in the hippocampus or cerebral cortex after 10-day CSF1R inhibition. The results are consistent with previous studies. A study has reported that 4 months of CSF1R inhibition commencing 1.5 months of age in 5xFAD mice did not alter Aβ levels or APP processing despite considerably depleting microglia (Spangenberg et al., 2019). Furthermore, a month-long CSF1R inhibition in an advanced stage of the disease (i.e., in 10-month-old 5xFAD mice) did not affect either Aβ plaques or soluble or insoluble fractions of Aβ (Spangenberg et al., 2016). Also, two-week CSF1R inhibition in aged 3xTg and APP/PS1 mice did not reduce Aβ plaques (Karaahmet et al., 2022). However, studies employing CSF1R inhibition for extended periods in 5xFAD mice reported diminished parenchymal plaque development (Spangenberg et al., 2019; Son et al., 2020). Thus, the overall impact of inhibiting CSF1R on Aβ plaques is minimal unless the inhibition is maintained for extended periods (Spangenberg et al., 2019; Son et al., 2020). However, inhibiting CSF1R for short or moderate durations can reduce synapse loss and neurodegeneration and increase circuitry connectivity in AD models (Spangenberg et al., 2016; Liu et al., 2021).

## Conclusions

Inhibition of CSF1R for 10 days during the early stages of neuroinflammation in 5xFAD mice resulted in the predominant removal of activated microglia in the hippocampus and cerebral cortex, leading to the remaining microglia displaying a non-inflammatory phenotype with decreased NLRP3 inflammasome activation. Such changes led to a substantially reduced proinflammatory microenvironment, decreased mTOR signaling, and increased autophagy. These findings suggest that periodically depleting activated microglia via short-term CSF1R inhibition following the onset of AD could be a promising approach to maintain a less proinflammatory environment, reduce mTOR activation, and enhance autophagy for extended periods in the AD brain. Such an approach may preserve better cognitive and mood function for extended periods.

## Acknowledgments

This work was supported by grants from the National Institutes of Health (National Institute for Aging grants, RF1AG074256-01 and R01AG075440-01 to AKS).

## Data availability statement

All data needed to evaluate the reported findings are present in the article.

## Author contributions

Concept: AKS. Research design and interpretation: AKS, MK, and LNM. Data collection and analysis: MK, LNM, YS, SA, CH, PKP, GS, SR, BS, JJG, CO, CH, RSB, SK, and AKS. The first draft of the manuscript text and figures: MK, LNM, YS, and SA. Finalization of manuscript text and figures: AKS, MK, and LNM. All authors provided feedback and approved the final version of the manuscript.

