## Supplemental File for "Residual Microglia Following Short-term PLX5622 Treatment in 5xFAD Mice Exhibit Diminished NLRP3 Inflammasome and mTOR Signaling, and Enhanced Autophagy"

**Supplemental Figure 1**

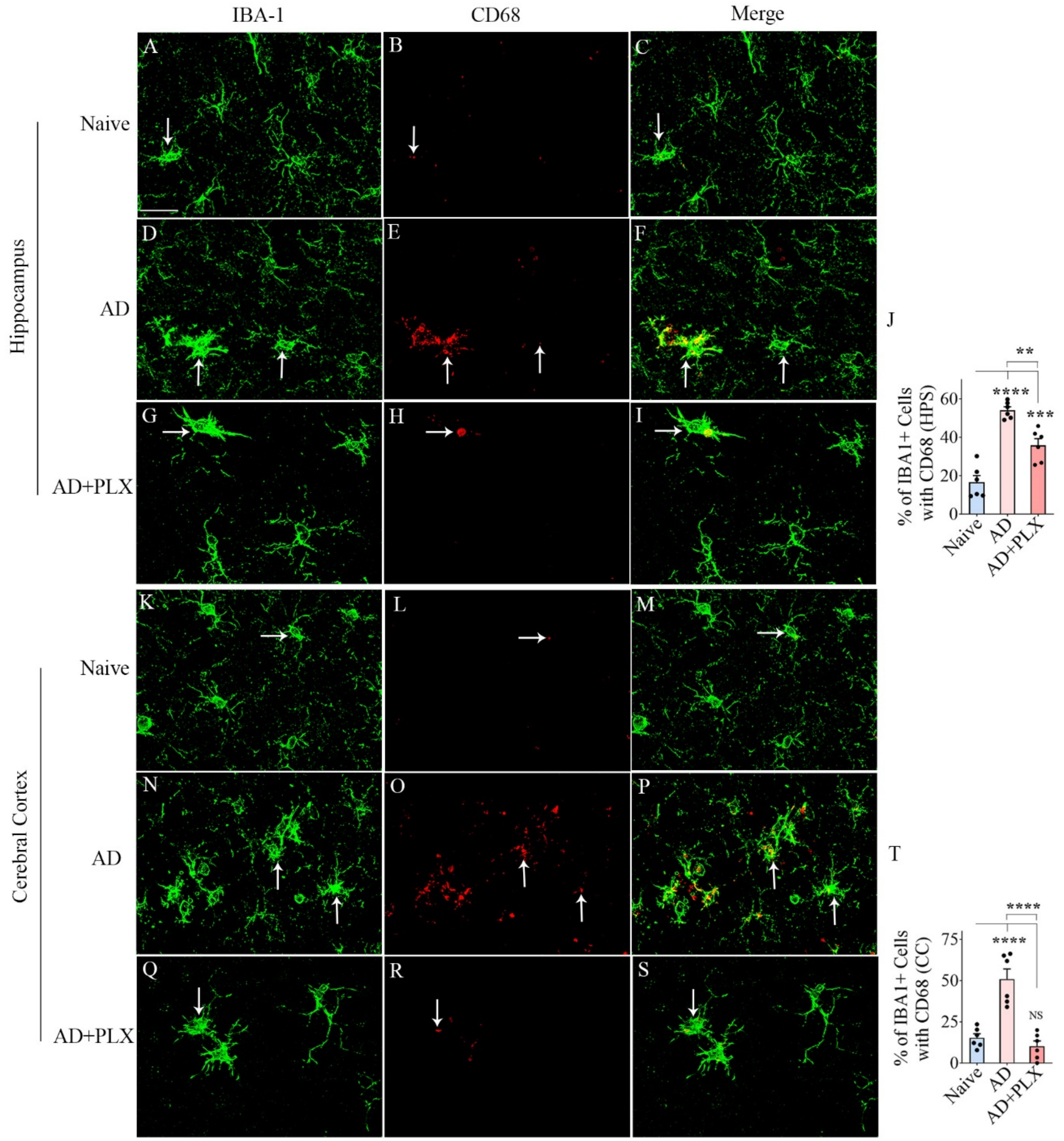


**Supplemental Figure 1:** ***Ten days of CSF1R inhibition in 3-month-old 5xFAD mice led to a reduced percentage of microglia expressing CD68 in the hippocampus and cerebral cortex.*** Figures A–I illustrate IBA-1+ microglia (green) displaying CD68+ structures (red) from the hippocampal CA3 subfield of naive (A-C), AD (D-F), and AD+PLX (G-I) groups. Figures K-S illustrate IBA-1+ microglia (green) displaying CD68+ structures (red) from the cerebral cortex of naive (K-M), AD (N-P), and AD+PLX (Q-S) groups. The bar charts in J and T compare the percentages of IBA-1+ microglia with CD68 in the hippocampus (J) and cerebral cortex (T). Scale bar, A-R = 10 µm; *, $p < 0.05$; **, $p < 0.01$; ***, $p < 0.001$; ****, $p < 0.0001$; NS, not significant.

**Supplemental Figure 2**

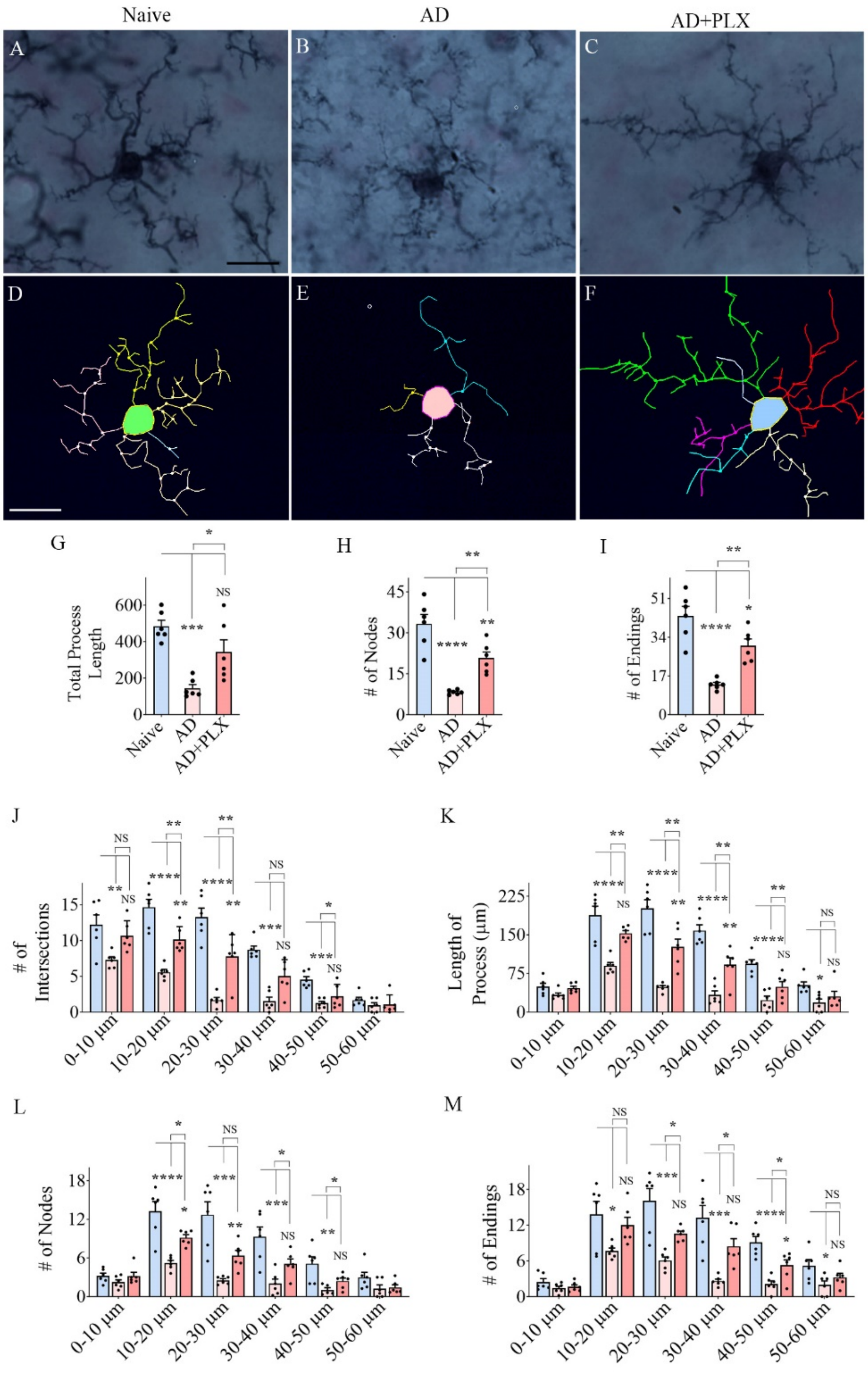


**Supplemental Figure 2: Residual microglia following *CSF1R inhibition displayed highly branching and ramified processes* in the cerebral cortex of *5xFAD mice.*** A-F shows representative examples of microglial morphology traced with Neurolucida from the cerebral cortex of naïve (A, D), AD (B, E), and

AD+PLX (C, F) groups. The bar charts G-I compare the various morphometric measures of microglia between naïve, AD and AD+PLX groups, which include the total process length (G), the number of nodes (H), and the number of process endings (I). The bar charts J-M compare the number of intersections (J), total process length (K), the number of nodes (L), and the number of process endings (M) between naïve, AD and AD+PLX groups at 0–10 μm, 10–20 μm, 20–30 μm, 30–40 μm, 40–50 μm, and 50–60 μm distances from the soma. Scale bar, A-F = 12.5 μm; *, $p < 0.05$; **, $p < 0.01$; ***, $p < 0.001$; ****, $p<0.0001$ NS, not significant.
